# Prolonged inflammation leads to ongoing damage after spinal cord injury

**DOI:** 10.1101/865089

**Authors:** Jacek M. Kwiecien, Wojciech Dabrowski, Beata Dąbrowska-Bouta, Grzegorz Sulkowski, Wendy Oakden, Christian J. Kwiecien-Delaney, Jordan R. Yaron, Liqiang Zhang, Lauren Schutz, Barbara Marzec-Kotarska, Greg J. Stanisz, John P. Karis, Lidia Struzynska, Alexandra R. Lucas

## Abstract

The pathogenesis of spinal cord injury (SCI) remains poorly understood and treatment remains limited. Emerging evidence indicates the severity of post-SCI inflammation and an ongoing controversy in the roles of astrocytes with studies identifying astrocytes as associated both with ongoing inflammation and damage as well as potentially having a protective role. We have completed an extensive systematic study with MRI, histopathology, proteomics and ELISA analyses designed to further define the severe protracted and damaging inflammation after SCI in a rat model. We have identified 3 distinct phases of SCI: acute (first 2 days), inflammatory (starting day 3) and resolution (>3 months) in 16 weeks follow up. Actively phagocytizing, CD68^+^/CD163^-^ macrophages infiltrate myelin-rich necrotic areas converting them into cavities of injury (COI) when deep in the spinal cord. Alternatively, superficial SCI areas are infiltrated by granulomatous tissue, or arachnoiditis where glial cells are obliterated. In the COI, CD68+/CD163^-^ macrophage numbers reach a maximum in the first 4 weeks and then decline. Myelin phagocytosis is present at 16 weeks indicating ongoing inflammatory damage. The COI and arachnoiditis are defined by a wall of progressively hypertrophied astrocytes. MR imaging indicates persistent spinal cord edema that is linked to the severity of inflammation. Microhemorrhages in the spinal cord around the lesion are eliminated, presumably by reactive astrocytes within the first week post-injury. Acutely increased levels of TNF-α, IL-1β, IFN-γ and other pro-inflammatory cytokines, chemokines and proteases decrease and anti-inflammatory cytokines increase in later phases. In this study we elucidated a number of fundamental mechanisms in pathogenesis of SCI and have demonstrated a close association between progressive astrogliosis and reduction in the severity of inflammation.

## INTRODUCTION

Spinal cord injury (SCI) is a leading cause of long term morbidity after motor vehicle accidents or trauma in combat, with no clinically standardized or substantially effective treatment. Effective therapeutics have remained limited due to an incomplete understanding of the pathogenesis of SCI and thus identified effective therapeutic targets while the underlying and persistent inflammatory damage after SCI remains poorly characterized. The role of invasive inflammatory cells and specifically astrogliosis in the ongoing damage after SCI is not fully defined with studies supporting both a role in SCI damage [1, 2] as well as a proposed potential protective role for astrocytes [3]. We have completed an extensive analysis with MRI, histochemistry, proteomics and ELISA in a rat model of SCI in order to further define the pathological changes that occur after SCI.

SCI initiates hemorrhage and ischemia, free radical release, severe inflammation, and cellular necrosis[4–6]. Pro-inflammatory cytokines such as interleukins 1β (IL1β) and 6 (IL-6) as well as tumor necrosis factor alpha (TNF-α) and interferon gamma (IFN-γ)[6–8], CC, CXC and CX3C chemokines [9] and matrix metalloproteinase 8 [10] are reported to be elevated after SCI, actively promoting severe inflammation.

After the initial injury there is an intense macrophage infiltration into damaged and necrotic areas, coinciding with conversion of damaged sites into a cavity of injury (COI), which blocks neuronal regrowth [11–14]. Axonal regrowth is inhibited by scar [2] and the fluid-filled spaces in the COI lesions [15]. The main characteristics of inflammation in SCI are ongoing spinal cord destruction, extraordinarily long duration and conversion of necrotic areas into the COI [12, 13] or arachnoiditis [16]. These changes after SCI are attributed to unique spinal tissue reactions, not observed in extra-neural trauma. Macrophage invasion into the COI and phagocytosis of myelin at 8 weeks post-SCI [17] indicate that tissue destruction continues over longer periods of time than is generally seen in the shorter duration of post-traumatic inflammation of extraneural tissues.

Damaged myelin has potent immunogenic functions in experimental allergic encephalomyelitis (EAE) [18] and after SCI [19] inducing severe lymphocyte-rich inflammation in EAE and macrophage-rich inflammation in SCI [12,13,17]. As previously reported in Long Evans Shaker (LES) rats lacking myelin [20–22] there is a reduction in inflammation after SCI [15]. In normally myelinated spinal cord, massive myelin damage induces a vicious cycle of tissue destruction, wherein macrophage phagocytosis of myelin [12,17,22] increases immune cell responses and further white matter damage, in turn attracting more macrophage infiltration.

The longevity of the inflammatory phase seen after SCI indicates that 24-48 hrs of intravenous infusion of high dose anti-inflammatory treatments such as methylprednisolone succinate [23] may provide a partial explanation for why short term high dose steroid infusions are unlikely to be effective as has been demonstrated by a lack of efficacy in prior clinical trials for steroid infusions given early after SCI. High dose steroids additionally cause severe toxicity [1], indicating the need for novel anti-inflammatory compounds of low toxicity that can target central mediators of ongoing inflammation and injury, with potential for sustained long-term administration designed to reduce prolonged inflammation after SCI [13, 24].

Astrocytes are immunocompetent [25, 26] secreting pro-inflammatory cytokines [27] and chemokines [28], complicating their role in SCI neuropathology [2]. Astrocytes have been proposed as mediators of SCI inflammation and scarring [2] but have also conversely been proposed as having a protective role in walling off the expanding cystic areas after SCI [3,14,29–32]. Interestingly, increases in reactive astrocytes parallel the rise of anti-inflammatory activity in rat SCI models [17]. In this study astrogliosis as the spinal cord response to injury and to supervening severe inflammation was systematically studied over a period of 16 weeks, exceeding the longevity of previous systematic studies addressing post-SCI astrogliosis [2]. The severe, diffuse astrogliosis that develops in the adult LES dysmyelinated rat [15] is associated with a rapid and total elimination of inflammation, at 7 days post-SCI [15]. In the present study the progressive severity of astrogliosis surrounding the COI or arachnoiditis was associated with decline in macrophage numbers and in general decline in spinal cord levels of pro-inflammatory cytokines, cytokines and other factors. Concurrently, general increase in anti-inflammatory cytokines (IL-2, IL-4, IL-13) and fractalkine (CX3CL1) was observed post-SCI suggesting that an anti-inflammatory process does develop over time along progressive astrogliosis, the reactive response in the CNS. Reactive astrocytes are associated with promotion of neuronal survival [33] and regulation of neural stem cell differentiation [34] indicating an association with actions that restore and preserve homeostasis after SCI [3]. Thus, the pro- and anti-inflammatory roles of astrocytes in spinal cord injury remain incompletely defined.

We present here an extensive time-course analysis of SCI pathogenesis including inflammatory destructive changes that correlate with neuronal damage and also potential neuroprotective reactions. We have further confirmed prior studies on SCI in a rat model of compression injury demonstrating progressive astrogliosis beginning 1-2 days post-SCI through week 16. The numbers of macrophages peak at 1-4 weeks post-SCI, with their gradual decline by 12-16 weeks post-SCI, while astrogliosis progressively walls off the COI coinciding with a reduction in macrophage invasion. These findings suggest that persistent astrogliosis around the COI is associated with gradual decrease in pro-inflammatory and increase in anti-inflammatory cytokines. We would postulate that there is potential for astrocyte mediated anti-inflammatory and neuroprotective activity as previously reported [3].

## MATERIALS and METHODS

### Ethical considerations

The studies on experimental rats were approved by the Animal Research Ethics Board at McMaster University, Canada, according to the guidelines and regulations of the Canadian Council for Animal Care. The animal experiments were performed at the Central Animal Facility, McMaster University. Given the invasiveness of the SCI ethical endpoints were instituted; rats with lethargy, hypothermia <35°C and ruptured urinary bladder were excluded from the study and humanely euthanized (total N=5).

### Experimental surgeries

Male, 16 week old Long Evans (LE) rats weighing 370-410g, (study total, N=90) at the maximum of their body size, were used for neurosurgeries and 16 week old Long Evans Shaker (LES) rats (N=5) were used as positive controls in proteomic analyses and in the immunohistochemistry.

The rats were housed in conventional, disease-free conditions and fed with rat chow and water *ad libidum*. Rats were induced with 5% isoflurane in oxygen and maintained under 3-4% isoflurane, the skin on the back was prepared surgically, laminectomy and the epidural balloon crush was performed as described previously [35]. Briefly, dorsal laminectomy was performed on the T10 vertebrum and a 3Fogarty catheter inserted over the intact dura rostrally to place the balloon over the mid-thoracic spinal cord. The balloon was inflated with 20 microL of saline for 5 seconds, then deflated and carefully removed [35]. After the spinal muscles were closed with absorbable sutures and the skin with stainless skin staples, the rats were administered 5 mL saline subcutaneously and 4 mg ketoprofen (Anafen, Meriel Canada, Inc.) for analgesia given subcutaneously prior to recovery from anaesthesia and returned to the holding room when awake. The painkiller was injected once daily for 2 more days at the same dose.

### Post-operative care

Experimental rats with the SCI became paraplegic and were maintained for 1, 2, 3, 7, 14, 28, 56, 84 and 112 days (N= 60, n=5). A large proportion (90%) of rats developed paralysis of the urinary bladder and were manually voided 1-2 times per day. Rats with hemorrhagic cystitis were treated with daily intramuscular injection of 50 microL enrofloxacin (50 mg/mL, Baytril®) for 3-5 days until blood cleared from urine. The bladder function was typically re-established within the second week post-SCI. Micturition was frequently observed in rats with distended bladder. The motor deficits gradually improved bilaterally or unilaterally to reach a permanent, unchanging deficiency by the 3^rd^ week post-SCI. The hind end paralysis coincided with rapidly progressive muscular atrophy of hind limbs and also with loss of body weight within the first 2 weeks. The body weight however, recovered partially in longer surviving rats.

### Whole body perfusion and tissue collection

The rats (n=5 for each post-SCI survival group and intact controls, N=60) were deeply anaesthetized with overdose of 80 mg sodium pentobarbital/kg body weight [35]. The chest was opened, blood was collected by the cardiac puncture into coagulating glass vials, 100 units heparin sodium then injected into the left ventricle and a cannula with flowing lactated Ringer’s solution inserted into the left ventricle. After blood was washed out, the flow was switched to formalin.

The spine including the spinal cord was removed, post-fixed in formalin and maintained in formalin in 50 mL plastic Falcon tubes for magnetic resonance imaging (MRI) studies. The serum was separated from coagulated blood samples, by centrifuge 45 min after the collection and stored at −85°C until assayed.

For proteomic and Western blots, the LE rats at 2, 7, 28 and 112 days post-SCI, intact negative controls (total N=20, n=5) and intact LES rats (N=5), were perfused with chilled, 4°C, lactated Ringer’s solution and the entire spinal cord removed promptly and snap frozen on dry ice. The spinal cords were stored at −85°C until assayed.

### MR imaging of spines

MR cross section images were taken from intact spines and at 1, 2, 3, 7, 14, 28, 84 and 112 days post balloon crush SCI. Prior to imaging, formalin-fixed spines were placed in a sealed plastic bag to prevent dehydration while allowing the imaging coil to be placed as close to the spine as possible. MRI experiments were performed on a 7 Tesla horizontal bore magnet (BioSpec 70/30, Bruker Biospin, Germany) using a quadrature rat brain receive coil (Bruker Biospin, Germany). T2-weighted RARE images were acquired with TR 3.6s, echo train length of 10, echo spacing of 8.6ms, effective TE 43ms, 12 signal averages, in-plane resolution of 0.1×0.1mm^2^, and 1mm slice thickness. In order to cover the entire length of the spine, images were acquired in 3 segments, and the spine was repositioned relative to the receive coil between segments. MR images were analyzed for number of quadrants of cord with evidence of cavitation (0=none vs. 1-4 quadrants), presence or absence of abnormality in a remote portion of cord, number of cord quadrants with evidence of blood product (0=none vs. 1-4 quadrants), number of cord quadrants with loss of grey/white matter contrast (0=none vs 1-4 quadrants), and longitudinal extent of cord involvement measured across multiple cord cross sectional images described as length (0mm, 1-10mm, 11-20mm, >20mm).

### Histology

Formalin in the Falcon tubes with spines was replaced by fresh formalin supplemented with 4% EDTA, pH 7.2 and placed on a rotating shaker to achieve decalcification and thus softening of the spinal bones. The decalcifying solution was replaced 2x per week and the spines became thoroughly soft within 7-8 weeks. The spines including the spinal cord and vertebral bones with bone marrow were cut perpendicular to the long axis at 3 mm thick fragments with the rostral face placed down in the tissue block and processed for histology. Paraffin blocks were cut into 6 μm thick sections and stained with luxol fast blue and counterstained with haematoxylin and eosin (LFB+H&E). Sections of the spine with the spinal cord were analyzed under a light 50i Eclipse Nikon microscope by an experienced experimental neuropathologist and the representative digital microphotographs taken and archived for further analysis.

### Macrophage counts in the cavity of injury

From each rat, 3 light microscopic images stained with LFB+H&E of 3 separate sections of the spinal cord were taken with the 40x objective. The images measured 225×300 μm and included the periphery of the cavity of injury with adjacent spinal cord tissue taking 20% of the area of the image. Macrophages; large cells with a large oval nucleus and abundant vacuolated cytoplasm with blue granules of myelin and/or with red blood cells, were counted and the average for each rat (n=5) in the survival group was averaged and standard deviation calculated.

### Immunohistochemistry

For GFAP immunolabelling, unstained sections were cut, mounted on glass slides and labeled with 1:200 dilution rabbit polyclonal antibody against rat glial fibrillary acidic protein (GFAP; Proteintech #16825-1-AP) prepared in 5% BSA/TBST incubated overnight at 4 °C. Slides were washed with TBST and labeled with 1:500 dilution HRP-conjugated goat anti-rabbit IgG (Abcam #ab97051) prepared in 5% BSA/TBST and incubated for 1 hour at room temperature in the dark. Slides were washed and staining developed with ImmPACT DAB Peroxidase Substrate (Vector Labs #SK-4105) for 30 seconds. Slides were counterstained with hematoxylin, dehydrated and mounted with #1.5 coverglass using Cytoseal™ XYL (ThermoFisher Scientific). A serially sectioned negative control omitting primary antibody was included in each staining batch.

For dual anti-CD68/CD163 labelling, glass-mounted sections were de-waxed, blocked with 2.5% normal horse serum (Vector ImmPress) and 1:100 dilution of mouse anti-rat anti-CD68(ED1) antibody (Abcam) applied for 30 min followed by horse anti-mouse Polymer (rat absorbed) RTU (Vector ImmPress) for 30 min after which the brown color was developed with peroxidase block and Novolink DAB (Leica). A 1:3000 dilution of rabbit anti-CD163 antibody for 15 min and then anti-rabbit Polymer AP (Leica) for 20 min resulting in magenta-red labeling. The dual CD68/CD163-labeled slides were counterstained with hematoxylin and coverslipped.

### Proteomic analysis of the spinal cord

Determination of cytokines, chemokines and other inflammatory mediators/factors totaling 34, in spinal cord homogenates was carried out using a rat cytokine array, panel C2 (RayBiotech, Inc.). The cell lysis buffer provided with the kit was used to obtain tissue lysates during homogenization of entire spinal cord. After extraction, the samples were centrifuged and supernatant was retained for analysis. The remaining steps of the procedure were performed according to the manufacturer’s instructions. Briefly, RayBio Rat Cytokine Antibody Array membranes were blocked at room temperature with 2 ml of blocking buffer for 30 min, then covered completely with samples (100 μg of spinal cord tissue protein) and incubated overnight at 4°C with gentle rotation. The samples were decanted and washed three times with 2 ml of wash buffer I and then with 2 ml of wash buffer II at room temperature with gentle shaking (5 min per wash). The working solution of primary antibody was prepared by adding 2 ml of blocking buffer to the tube containing biotinylated antibody cocktail. The mixture was added to each membrane, and incubated overnight at 4°C with gentle rotation. After washing, 2 ml of 1000-fold diluted horse radish peroxidase– conjugated streptavidin was added to each membrane and incubated at room temperature for 2 hours. The samples were washed with wash buffer I and then with wash buffer II. Detection buffer A and B provided in the kit were mixed, pipetted onto the membranes, and incubated for 2 min. at room temperature. Signals were detected directly from the membrane using a chemiluminescence imaging system. Comparison of the signals with the table included in the kit enabled identification of released cytokines. A relative analysis of the intensity of the dots was performed with ImageQuant TL software, with comparison to negative, non-injured LE rats.

### Western blot analysis of the spinal cord

Proteins of spinal cord homogenates were separated by SDS-PAGE and then electro-transferred onto nitrocellulose membranes. The membranes were incubated in PBS (80mM Na_2_HPO_4_, 20mM NaH_2_PO_4_, 100mM NaCl; pH 7.5) containing 0.05% Tween 20 (PBS-T) and 5% milk powder for 1.5 h at room temperature. The membranes were then washed in PBS-T and incubated overnight at 4°C in PBS-T containing 5% milk powder and primary antibody: monoclonal anti-GFAP (Sigma-Aldrich, G3893) in dilution 1:400 or monoclonal anti-actin (MP Biomedicals, 0869100) in dilution 1:500 (internal standard). After washing in PBS-T, membranes were incubated for 30 min at room temperature in PBS-T containing 5% milk powder and secondary antibody: peroxidase-conjugated anti-mouse IgG (1:5000) (Sigma-Aldrich, A2304). The membranes were then washed in PBS-T, the cross-reacted antibodies were detected using a chemiluminescence system (ECL Western Blotting System, Amersham Bioscences) and exposed to Hyperfilm ECL. The films were scanned and quantified using Image Quant TL v2005.

### ELISA of the serum

The ELISA kits for GFAP, myelin basic protein (MBP), and light chain of neurofilament (NF-L) (MyBioSource) were used to measure the levels of these proteins in the serum samples from the SCI rats according to the supplier. The selected markers represent the myelin (MBP), astrocytes (GFAP) and axons (NF-L) damaged by the trauma and then by the destructive inflammation as indicated by histologic analysis.

### Statistical analyses

Statistics for MRI, cell counts and proteomics were calculated by ordinary One-Way ANOVA with Tukey’s post-hoc compared against Day 1 post-SCI in GraphPad Prism v8.1.2. \**p*<0.05, \*\**p*<0.01, \*\*\**p*<0.001, \*\*\*\**p*<0.0001, n.s. is not significant.

## RESULTS

### T2 Magnetic resonance imagining (T2-MRI) after SCI

A longitudinal analysis of MR images from rat spinal cord after SCI demonstrates significant changes in local damage, hemorrhage and inflammation after SCI (Fig. 1). COI formation begins by day 7 and continues to increase in size (day 14, P<0.01: day 28, P<0.05) stabilizing at days 84 (P<0.0001) to 112 (P<0.01) (Fig. 1A). Areas of hemorrhage proximal to the site of the crush injury, and separated from it by undamaged spinal cord, are observed within the first 2 weeks post-SCI (days 1, 2, 7, P<0.01) (Figure 1B). Remote areas of hemorrhage have previously been observed and are considered related to increased vascular fragility and unrestricted movements of the rats (Fig 1B) [12]. Hemorrhagic areas at the site of SCI are seen early and then are reduced (P < 0.05 to 0.001 for days 1-112) (Fig. 1C). Loss or reduction in gray and white matter (G/W) border differentiation is representative of inflammation and edema [4] and is detected early, also persisting to day 112, (Fig. 1D, P<0.001). Overall extent of cord damage, caudal to distal, is seen in the first 14 days and continues up to day 84 post SCI (Fig 1E). MR images are representative at sites of SCI at individual follow up times in Fig. 1F, illustrating COI and hemorrhagic areas.

**Fig. 1.**
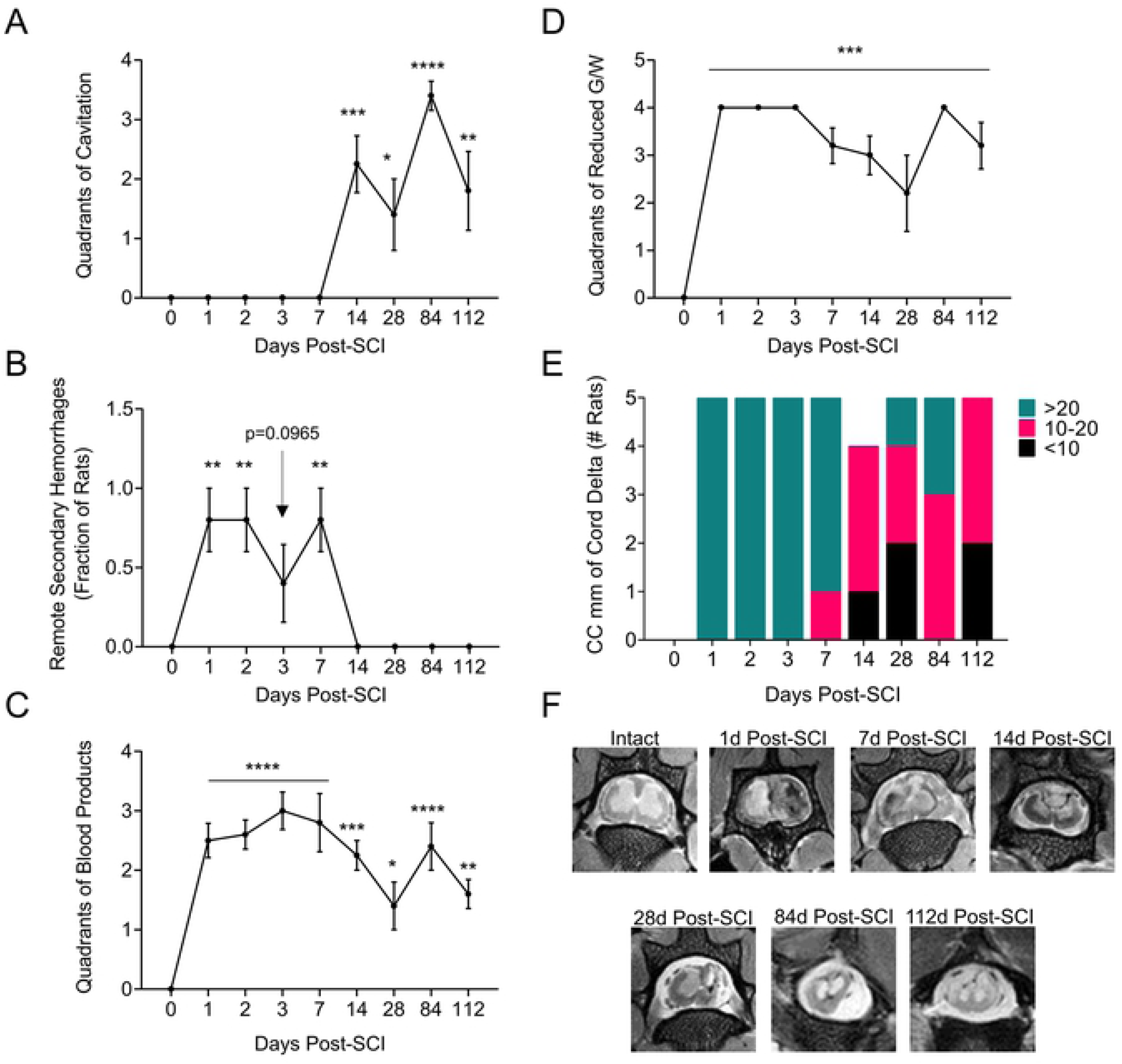
MRI analysis demonstrating disease progression. Cross sectional images recorded from spines, perfusion fixed *in situ* after euthanasia at each follow up time and analyzed blinded. A. Number of quadrants of cord with evidence of cavitation. 0=none vs. 1-4 quadrants. B. Presence or absence of abnormality in a remote portion of cord, numbers of rats with evidence of remote damage. C. Number of cord quadrants with evidence of blood product. 0=none vs. 1-4 quadrants. D. Number of cord quadrants with loss of grey/white contrast. 0=none vs. 1-4 quadrants. E. Longitudinal extent of cord involvement (measured across multiple cord cross sectional images described a s length. 0=none, 1-10mm, 11-20mm, >20mm F. MRI images – cross sections taken at days 0, -r90r to injury, and 1, 2, 7, 14, 28, 84 and 112 days post balloon crush SCI. Measured changes demonstrate statistically significant changes in cord damage. ANOVA analysis *P < .05, **P < 0.01, ***P < 0.001, ****P < 0.0001

### Histopathologic changes after SCI

A detailed comparison of MRI and histopathology of the rat SCI over 112 days defines three distinct phases of injury and repair; Phase 1 Acute, Phase 2 Inflammatory, Phase 3, Resolution (Fig. 2).

**Fig. 2a.**
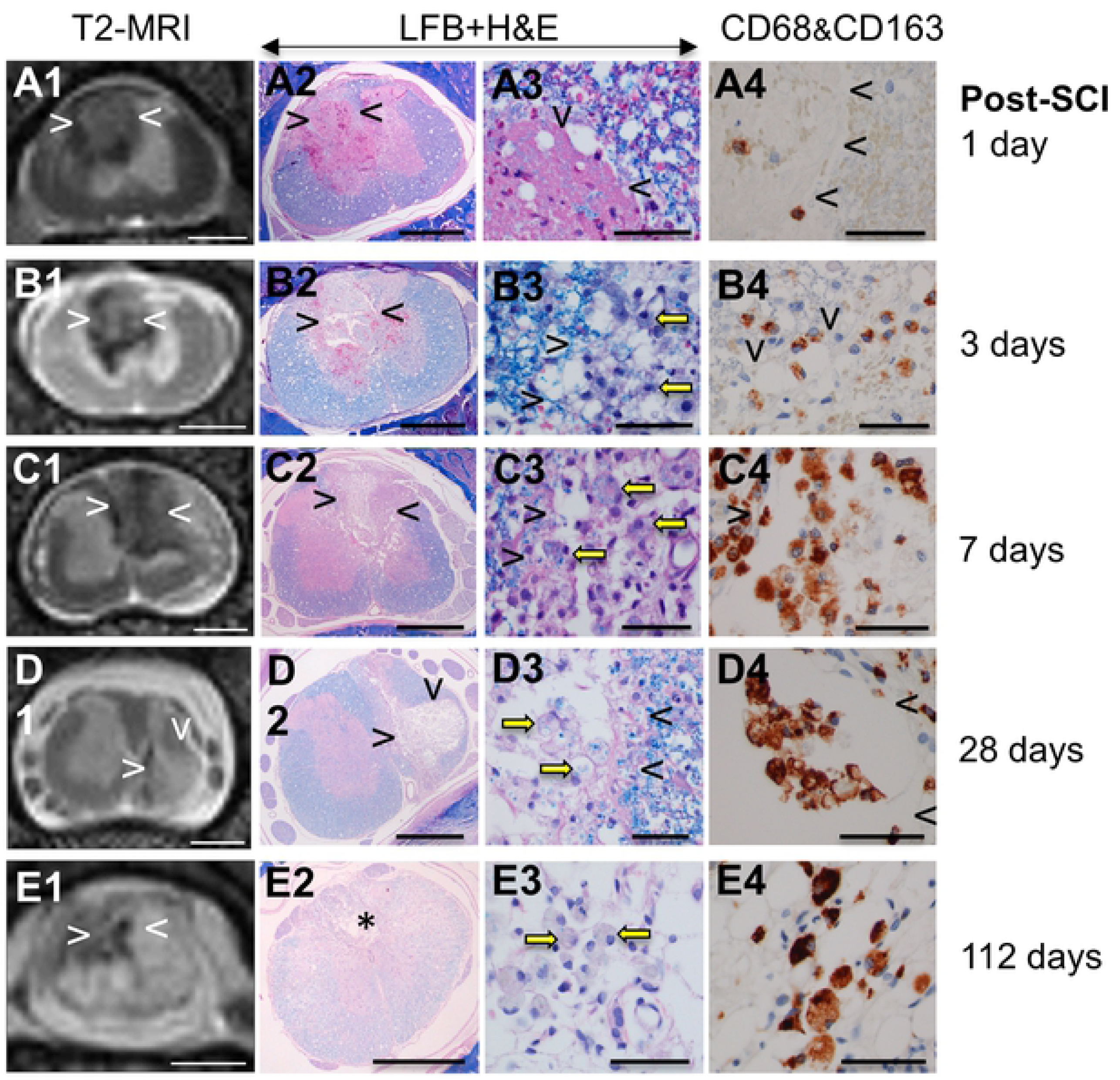
Spinal cord injury resolving in the cavity of injury. The T2-weighed MRI of the spinal cord injury (SCI, column 1) with histology of a low magnification (2x, column 2), and high magnification (60x, column 3), with dual anti-CD68 (brown color) and anti-CD163 (red magenta color) antibody staining immunohistochemistry (column 4) of the site of injury in rats at 1 day (A), 3 days (B), 7 days (C), 28 days (D,), and 112 days (E, F) post-SCI. The arrowheads indicate the cavity of injury which is also indicated by an asterix in E2. MRI and histology demonstrate acute intermediate and chronic phases of damage after SCI. Figures A1-E1, column 1 compares MRI to low power histology images B2-E2 column 2 at each follow up time, 1, 3, 7, 28 and 112days post SCI. Column 3 A3 to E3 demonstrates inflammatory cell invasion defined by luxol fast blue staining. Column 4 A4 to E4 illustrate M1 macrophage invasion at each follow up time. Size bars: A – 2 mm; B – 1 mm, C – 100 μm, D, E – 50 μm.

**Fig. 2b.**
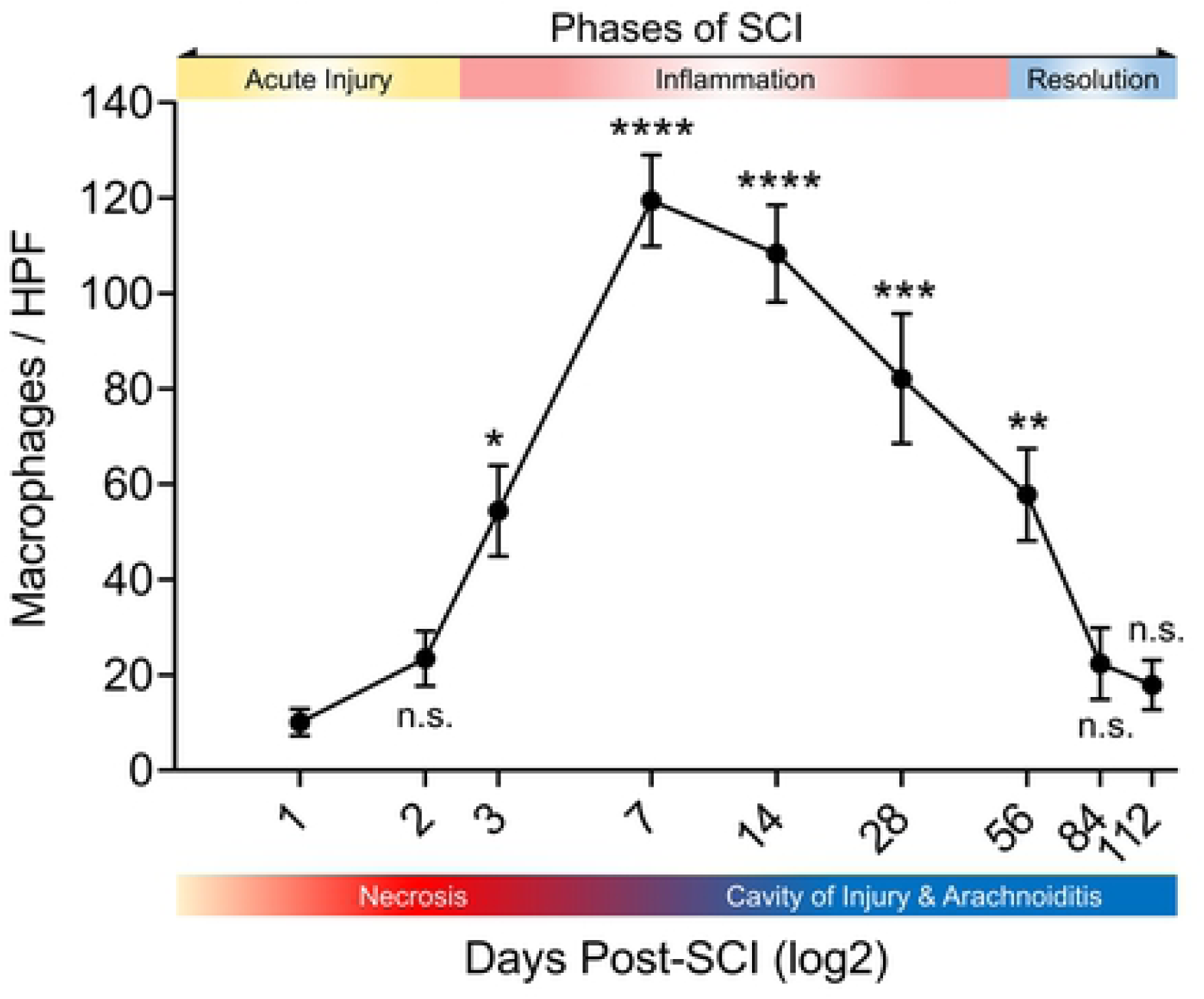
Macrophage counts in the cavity of injury along 3 phases of initiated by the spinal cord injury. One Way ANOVA with Tukey’s post-hoc compared against Day 1 post-SCI.

In the rat SCI the *acute phase* lasts for 2 days and is characterized by hemorrhage and necrosis (Fig. 2a.A1-A4). These areas are poorly defined on T2-weighted MRI. Increased intensity is believed due to reduced anisotropy when compared to intact areas of white matter (Fig. 2a.A1) or in intact control spinal cords (Fig. 1F). Necrotic areas are surrounded by a loss or reduced definition of the G/W border, while histologic definition is preserved (Fig. 2a.A2). Differences between the MRI and histology provide an indication of edema surrounding the trauma site, detected at this acute stage by T2-MRI, but not by histology. Extravasation of numerous scattered red blood cells (RBCs) (Fig. 2a.A3) indicates microvascular damage potentially causing vasogenic edema. Rare mononuclear cells detected in necrotic areas and in the surrounding spinal cord are CD68^+^/CD163^-^ (Fig. 2a.A4).

At day three post-SCI, two distinct inflammatory reactions develop depending on the location of the area of necrosis in the spinal cord and on the apparent influence of spinal cord tissue on the inflammatory process. Deep in the spinal cord, necrotic areas have increased intensity on T2 MR images indicating the accumulation of fluid with edema in surrounding spinal cord (Fig. 2a.B1,2). Numerous large cells, often containing intracytoplasmic granules of myelin or RBCs (Fig. 2a.B3), infiltrate the periphery of the injury site. A large proportion of these cells are CD68^+^/CD163^-^ (Fig. 2a.B3, B4) with rare cells scattered in the surrounding spinal cord. At day 7 post-SCI (Fig. 2a.C1-C4) a necrotic area deep in the spinal cord is seen as more intense in contrast to the surrounding tissue on T2-MRI, with more edema in the surrounding tissue indicated by blurring of the G/W border. This apparent fluid shift into the area of necrosis coincides with the creation of the *cavity of injury* (COI) (Fig. 2a.C,D). Large numbers of CD68^+^/CD163^-^ M1 pro-inflammatory macrophages, often containing myelin granules and/or RBCs in the COI and in the surrounding spinal cord are seen at 1-4 weeks post-SCI (Fig. 2a.C3,C4,D3,D4). The necrotic debris, including damaged myelin and RBCs is removed completely from the COI by day 14 (not shown). In the subsequent weeks, numbers of macrophages gradually decline in the COI and in surrounding spinal cord. By week 16 (Fig. 2a.E1-E4) the COI is filled with clear fluid with pockets of CD68^+^/CD163^-^ macrophages with myelin granules (Fig. 2a.E3) indicating persistently active and destructive inflammation.

The histologic analysis of the bone marrow (Fig. 3b) indicates a peak of CD68^+^ cell hyperplasia at the day 7 post-SCI with reversal after 14 days to a balanced representation of CD68^+^ and/or CD163^+^ and CD68^+^/CD163^+^ cells post-SCI.

**Fig. 3a.**
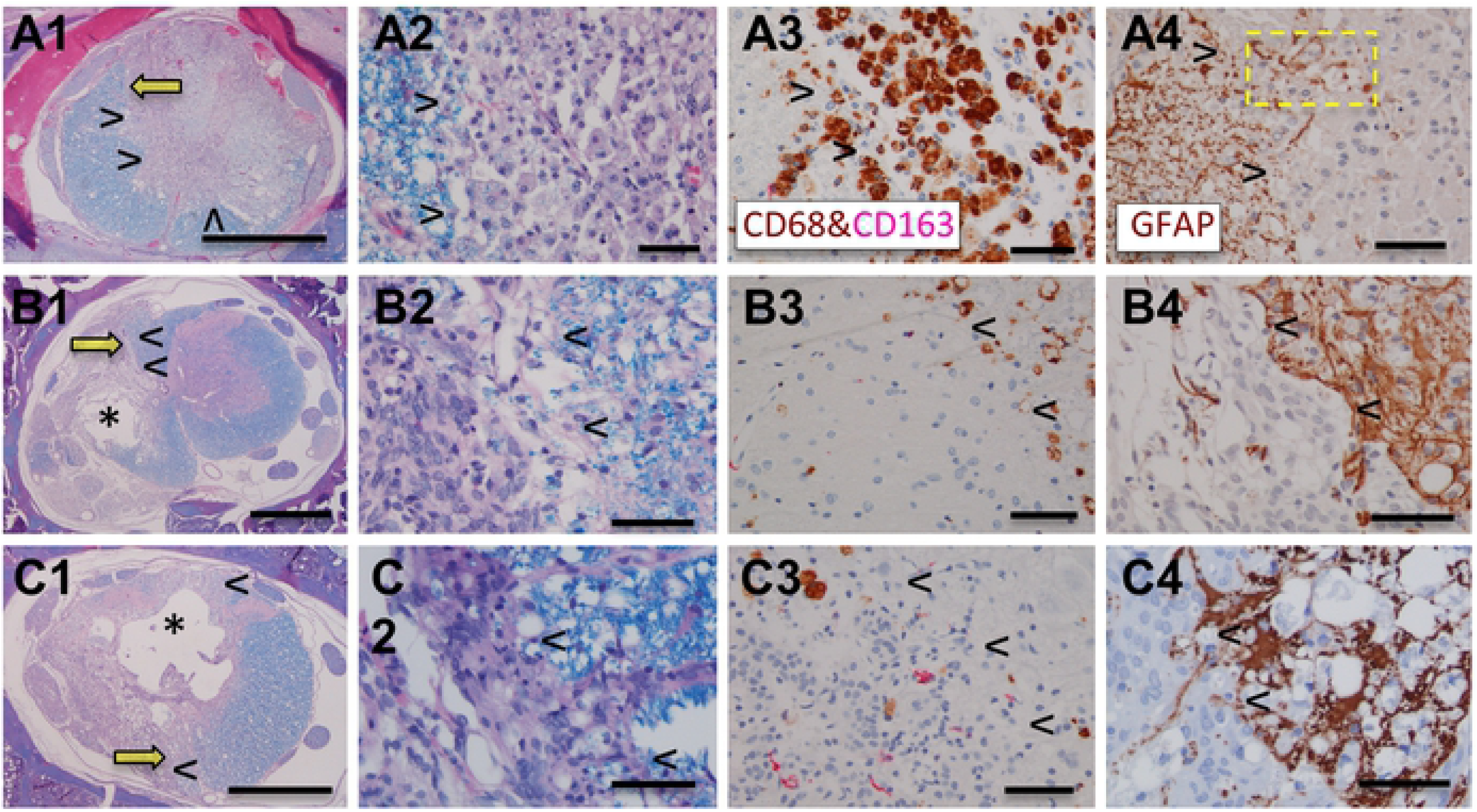
Spinal cord injury resolving in arachnoiditis. The columns 1 and 2 show large areas of obliteration of the spinal cord by arachnoiditis (arrowheads). Yellow arrows in column 1 indicate an area magnified in column 2. The asterix in B1, C1 indicates the cavity of injury (COI). The arrowheads in columns 3 and 4 indicate the margin between arachnoiditis and the spinal cord pointing towards arachnoiditis. The yellow box in A4 indicates astrocytic processes protruding into the area of arachnoiditis likely to be obliterated by severe infiltration by CD68^+^/CD163^-^macrophages (see A3). Luxol fast blue counterstained with hematoxylin and eosin (LFB+H&E); A1, A2, B1, B2, C1, C2. Dual immunohistochemical stain with antibodies against CD68 (brown) and against CD163 (magenta); A3, B3, C3. Anti-GFAP antibody stain; A4, B4, C4. Size bars: A1, B1, C1 – 1,000 microns, A2-A4, B2-B4, C2-C4 – 50 microns.

**Fig. 3b.**
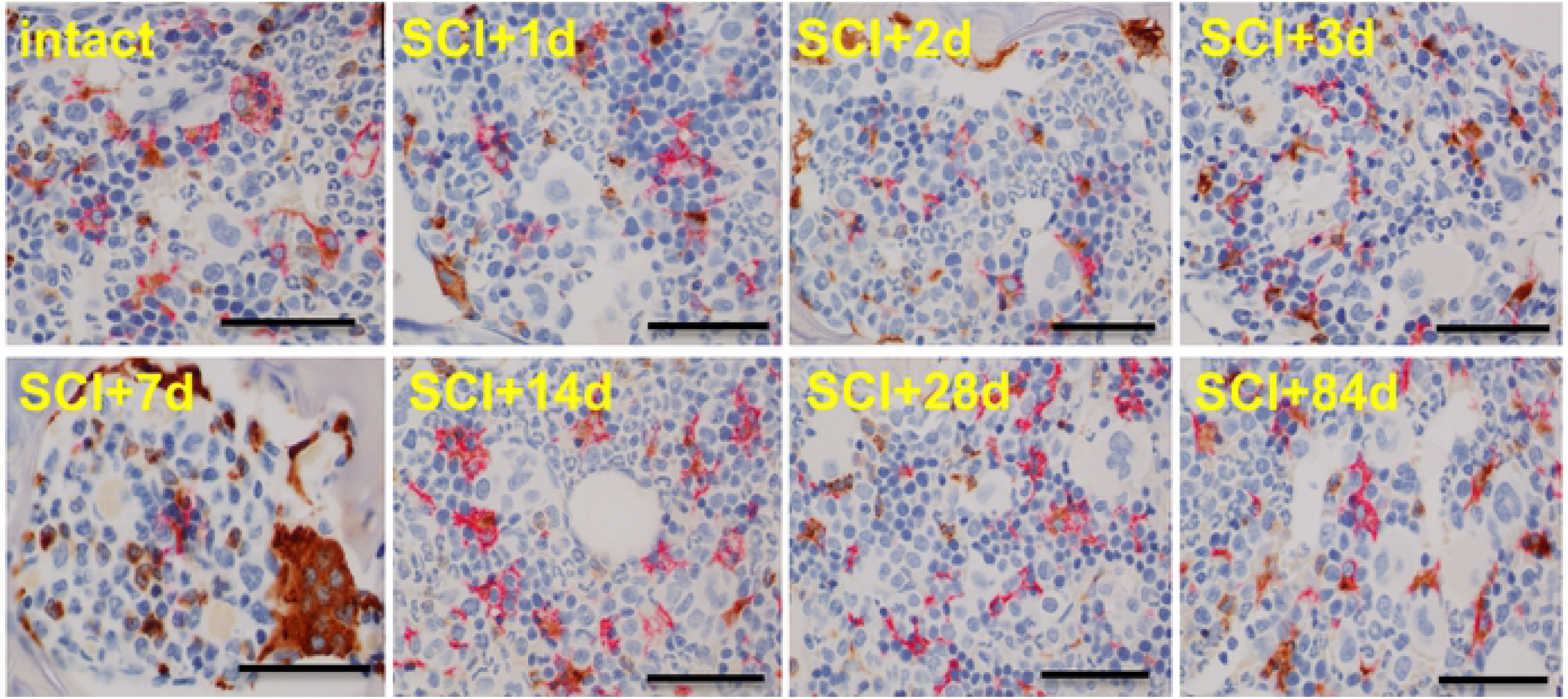
Bone marrow following the SCI. Dual anti-CD68 (brown color) and anti-CD163 (magenta color) labelling of cells of monocytic lineage in the vertebral bone marrow in the intact rat and in injured rats for up to 84 days post-SCI. At the day 7 post-SCI there is a remarkable hyperplasia of CD68^+^ cells. Size bars; 50 μm.

In areas where trauma-induced necrosis involved the periphery of the spinal cord there is an invasion of inflammatory cells from the subarachnoid space (Fig. 3a.A-C), within the 1^st^ week post-SCI. This inflammatory infiltration called *arachnoiditis* [16] - involves macrophages laden with granules of myelin and/or red blood cells, fibroblasts and capillary blood vessels and has an appearance of a granulomatous infiltration observed in traumatic injuries of soft, extra-neural tissues. Arachnoiditis appears to expand into the spinal cord with its concurrent obliteration as indicated by scattered fragments of GFAP-positive processes among inflammatory cells (Fig. 3a.A4, B4, C4). The intensity of infiltration of CD68^+^/CD163^-^ macrophages in arachnoiditis is severe starting at 7 days post-SCI but then declines by 4 weeks post-SCI while CD68^+^ and CD163^+^ macrophages are rare and scattered by 12-16 weeks post-SCI (Fig. 3a.C3). Ultimately, arachnoiditis becomes a scar tissue that is devoid of astrocytes (Fig. 3a.A4, B4, C4), but consistently increasing numbers of GFAP^+^, remarkably enlarged astrocytes in the adjacent spinal cord (Fig. 3a.A3, B3, C3) indicate a progressive astrogliosis, as a barrier against the severity of granulomatous inflammation.

At 3-4 months post-SCI the numbers of macrophages in the COI decline to a plateau (Fig. 2b). These macrophages still contain granules of myelin (Fig. 2a.E3) and they are CD68^+^/CD163^-^ (Fig. 2a.E4), pro-inflammatory, M1 type. However, their low number and persistence in isolated small pockets also suggest a declining inflammation probably due to an anti-inflammatory activity of the spinal cord that is water-soluble with its potency gradually increasing as the numbers of macrophages decline. Arachnoiditis changed morphologically from an inflammatory granulomatous tissue rich in infiltrating CD68^+^/CD163^-^ macrophages into a mature scar with few scattered CD68^+^ and CD163^+^ macrophages (Fig. 3a.A1-A3, B1-B3, C1-C3).

### Astrogliosis post-SCI

Astrocytic reaction in areas of SCI is associated with hypertrophy of GFAP^+^ cellular processes and their re-alignment in the spinal cord initially surrounding the area of necrosis and later the COI (Fig. 4.A). At 2-4 weeks astrocytic processes form a discontinuous and at 8-16 weeks post-SCI a continuous thick membrane facing the liquid content of the COI which coincides with the increase in numbers of hypertrophic astrocytes in the belt approximately 100-200 microns thick, in the surrounding spinal cord (Fig. 4.A). In a similar anisomorphic reaction, astrocytic hypertrophy and hyperplasia develop gradually from 1-16 weeks post-SCI in the spinal cord facing arachnoiditis (Fig. 3a.A4, B4, C4).

**Fig. 4.**
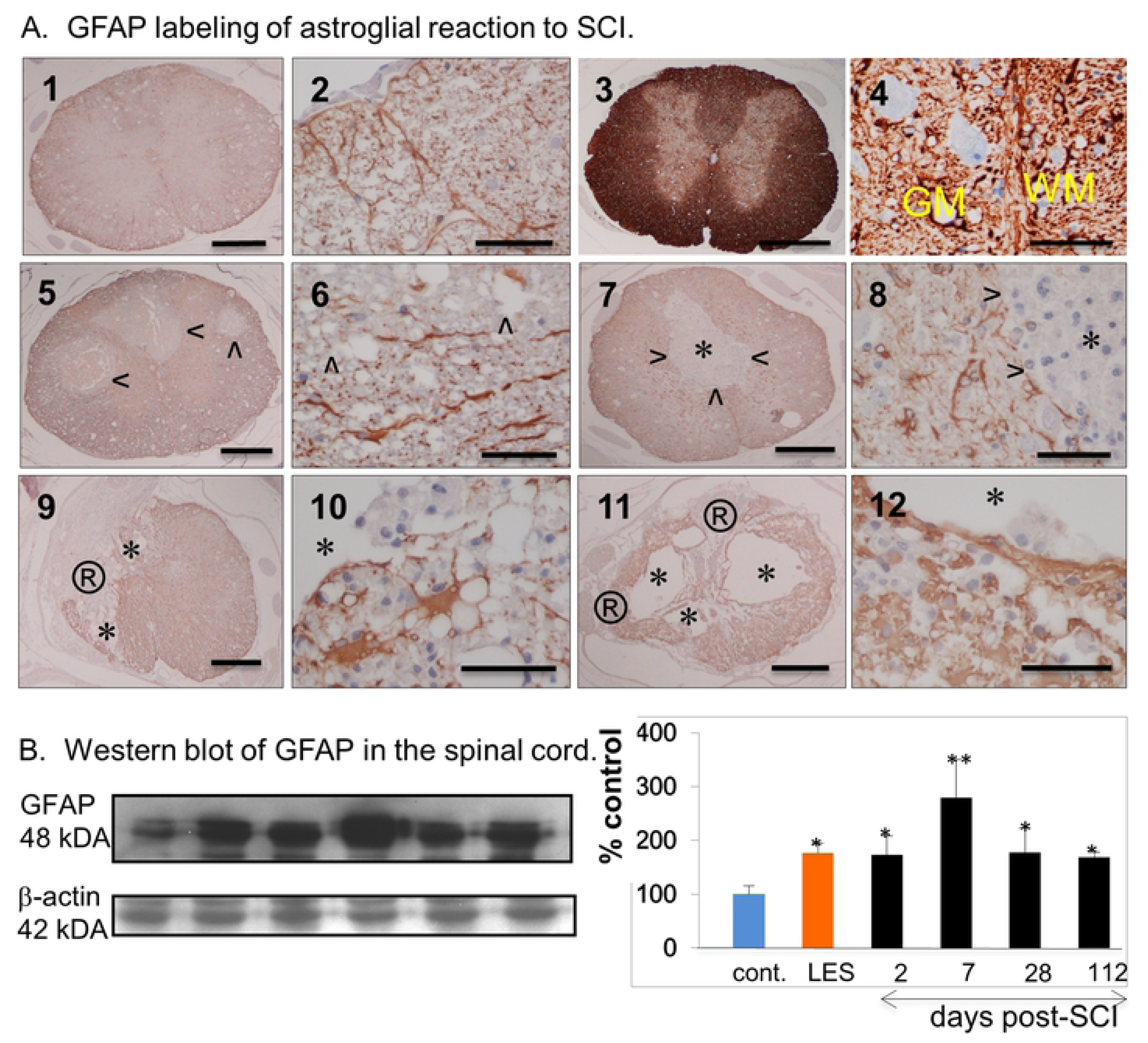
Astrogliosis in the SCI. A. Anti-GFAP antibody labeling of intact LE rat (1,2), intact LES rat (3,4) and post-SCI rat spinal cord at 2 (5,6), 7 (7,8), 28 (9,10) and 112 days (11,12). Arrowheads indicate the areas of injury that are GFAP^-^, asterix indicates a COI, ® indicates an area of arachnoiditis. Yellow lettering in (4) indicates gray matter (GM) and white matter (WM). Size bars; 1,000 microns -1, 3, 5, 7, 9, 11; 50 microns – 2, 4, 6, 8, 10, 12. B. Western blot of GFAP in the spinal cord. The Western blots from the protein extracted from the spinal cord of intact LE-cont. rats, intact LES rats and of injured rats at 2, 7, 28 and 112 days post-SCI are shown on the left and the densimetry results expressed as the percent of the negative control or intact LE-control rat.

### Astrocytic erythrophagocytosis

Although the ingestion of extravascular RBCs by macrophages (erythrophagocytosis) in the necrosis and then in the COI is abundant in the first 2 weeks post-SCI, numerous RBCs from micro-hemorrhages scattered in the surrounding spinal cord, are apparently removed by a different mechanism (Fig. 5a). While at 3 days post-SCI, large stellate cells encompassing multiple RBCs are numerous and morphologically compatible with astrocytes, CD68^+^ macrophages are relatively rare and small in the spinal cord while more numerous and large, often encompassing multiple RBCs in the site of necrosis.

**Fig. 5a.**
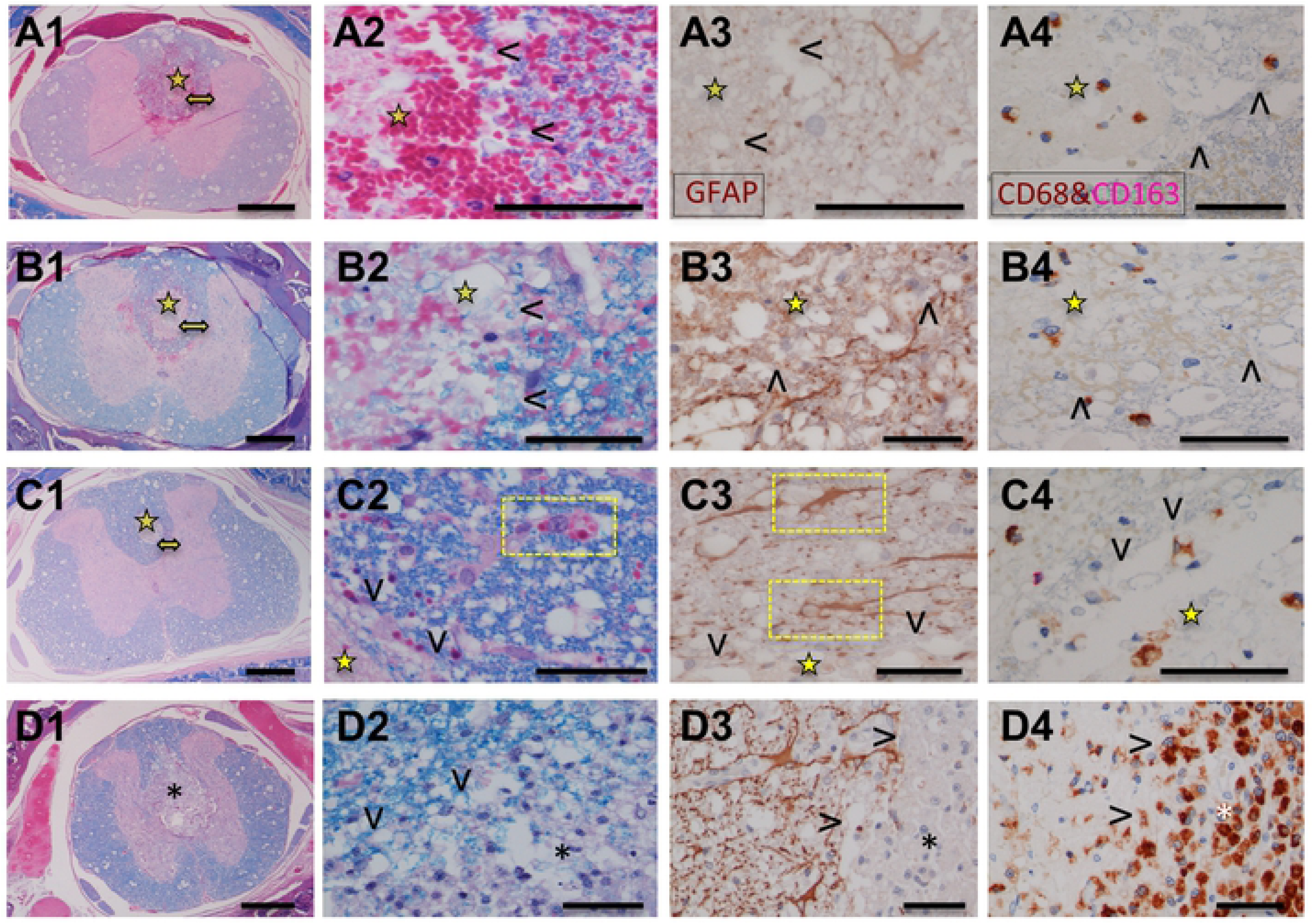
Astrocytic erythrophagocytosis following the SCI. Columns 1 and 2 present histology at 1, 2, 3 and 7 days post-SCI (A-D respectively). The column 3 have sections labelled with anti-GFAP antibody and column 4 labelled with anti-CD68 (brown) and anti-CD163 (red magenta) antibodies. The arrowheads (columns 2-4) indicate the area of necrosis and the asterix (row D) indicates the COI infiltrated by macrophages. The yellow star indicates the area of necrosis and double headed yellow arrow in column 1 indicates peri-lesional area magnified in column 2. The yellow boxes indicate internalization of red blood cells by astrocytes. Size bars; 1,000 microns in the column 1; 50 microns in the column 2-4.

### Ependymal cell plasticity

In the intact central canal of the spinal cord, GFAP^-^ ependymal cells form a pseudo-columnar epithelium with the brush border facing the lumen (Fig. 5b). At 7 days post-SCI and with inflammatory infiltration nearby numbers of ependymal cells increase and become GFAP^+^ extending their processes away from the central canal. At 2 weeks post-SCI clusters of GFAP^+^ cells are connected to the hypertrophied ependyma of the central canal. At 16 weeks post-SCI, the area of the central canal adjacent to COI and arachnoiditis is populated by scattered isolated or adjoining, clusters of epithelial cells often forming rosettes. Only a proportion of these cells are GFAP^+^, the remaining ones stain faintly positive or negative.

**Fig. 5b.**
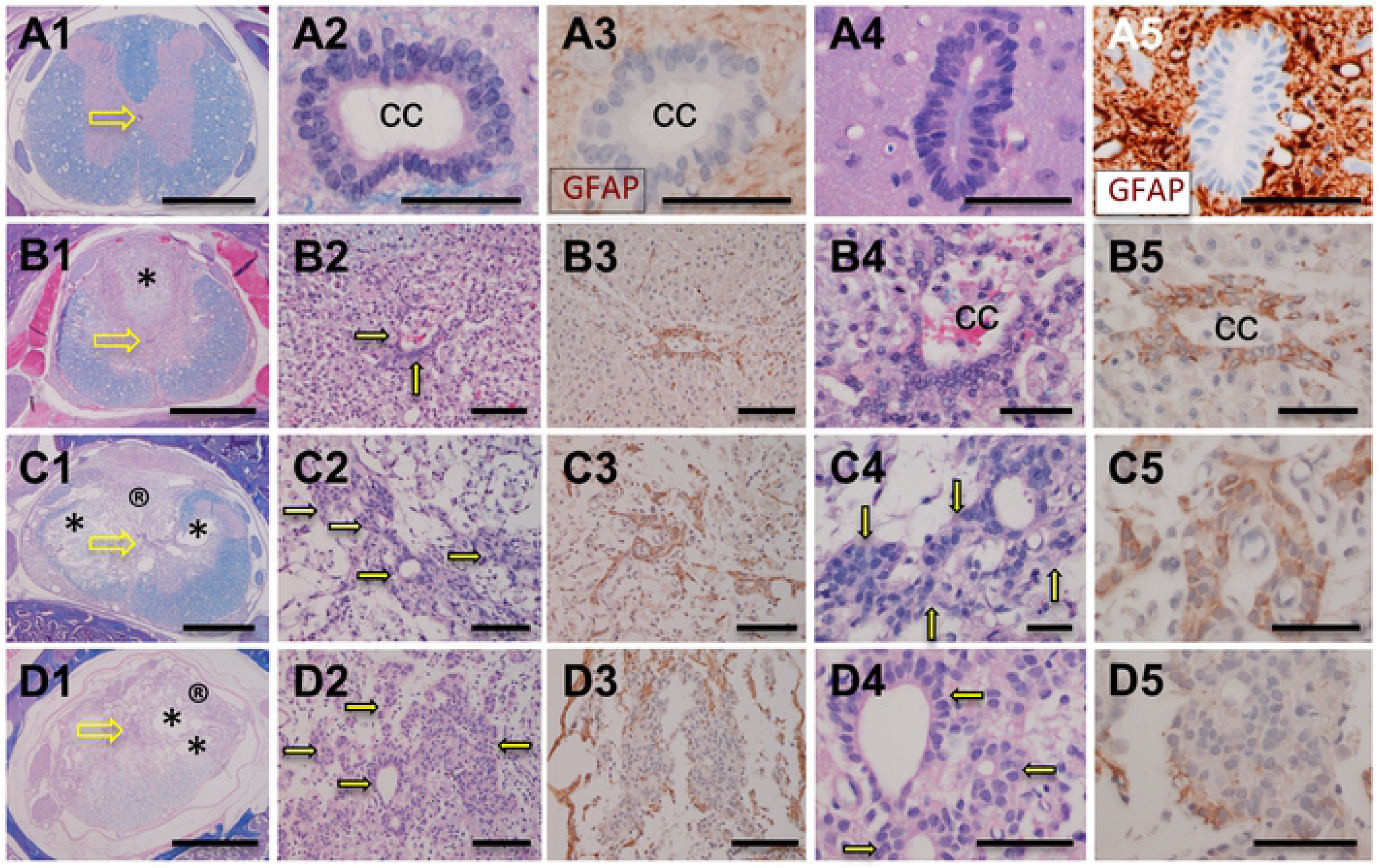
Ependymal cell plasticity after spinal cord injury. Histology of ependymal cells of the central canal (cc) in intact LE (A1-3), intact LES (A4,5) and in the spinal cord of LE rat at 7 (row B), 14, (row C) and 112 days (row D) post-SCI. Sections in the columns 1, 2, 4 are stained with luxol fast blue and counterstained with haematoxylin and eosin (LFB+H&E). Columns 3 and 5 are stained for GFAP. In the column 1, yellow empty arrows indicate the location of the central canal. The solid yellow arrows indicate clusters of epithelial cells interpreted as ependymal cells. Size bars: column 1 – 1,000 microns; columns 2, 3 – 100 microns; columns 4, 5 – 50 microns.

### Proteomic analysis of the spinal cord tissue post-SCI

A proteomic analysis of 34 factors involved in neuroinflammation was performed on protein extracts from the entire spinal cord collected at 2, 7, 28 and 112 days post-SCI. Changes in levels of each proteomic marker were measured and are provided as a percentage change when compared to the negative LE controls (e.g. intact spinal cords) (Fig. 6). The spinal cord of the LES rat is included as a positive control because of a previously demonstrated anti-inflammatory activity [15] and, preserved neuroplasticity in adult rats [15, 36]. The results of the analysis are presented as the mean percent of the detected levels of individual factors vs intact LE rats and analyzed with two way ANOVA (Fig. 6).

**Fig. 6.**
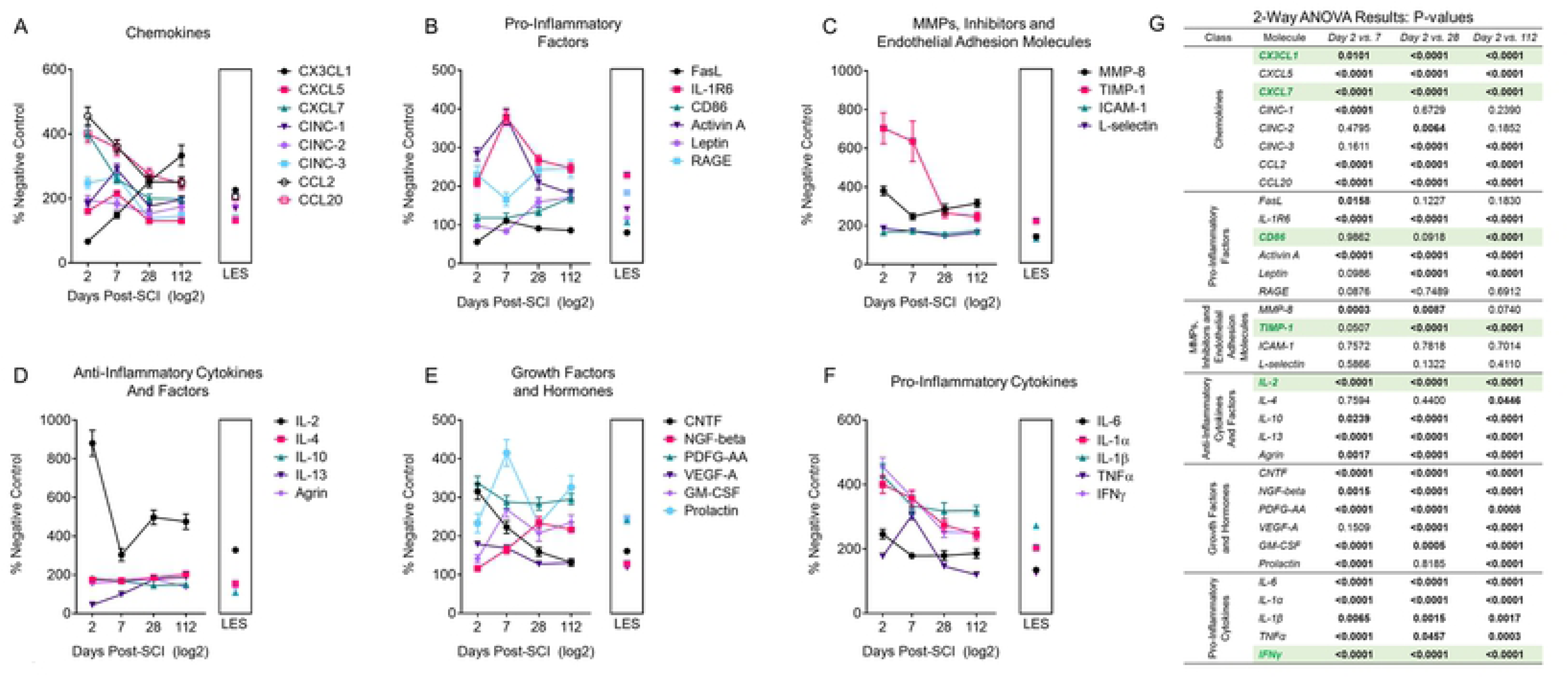
Dynamic changes in levels of 34 factors in the spinal cord following the SCI. The levels of factors are expressed as the percent of intact negative control, LE-cont. rat = 100%. The values obtained from the intact LES rat serving as the positive control are shown in the vicinity of the results from the rats sampled at 112 days post-SCI in the box. Levels of each of 34 factors were compared between 2 and 7 days, 2 and 28 and 2 and 112 days post-SCI, analyzed by 2-way ANOVA and p values presented in the adjacent table.

Levels of pro-inflammatory cytokines including IFN-γ, TNF-α, IL1β, IL-6, matrix metalloproteinase-8 and chemokines are increased within the first week post-SCI. Select factors had a demonstrated subsequent decline (TNFa, CCL7, CXCL5,7 and others demonstrated an ongoing increased expression at values similar to these of the positive control (LES rat, interleukins, CX3CL1). The high levels of IL-2, 900% of intact control at 2 days post-SCI remain markedly elevated for the entire duration of the study which contrasted with IL-4, IL-10 and again whose levels were only moderately elevated, <200% intact control. Il-13 is remarkably reduced within the first week post-SCI then elevated to approximately 200% of intact control. Overall there is a persistent and marked elevation of cytokines and chemokines, pro-inflammatory factors throughout the study with values ranging from 200 to 800% over control.

GFAP expression measured on Western blot is correlated with immunohistochemical changes associated with specific phases after the SCI (Fig. 4). Although GFAP expression increased by 200% of intact control on day 2 and by 300% on day 7 post-SCI, it declines to 200% on the day 28 and 112. Histology demonstrated a persistent increase in expression of GFAP indicating ongoing and progressive astrogliosis (Fig. 3a.A, Fig. 4.A4,B4,C4,).

### Biomarkers of the SCI in the serum

Changes in levels of 3 biomarkers; GFAP, MBP and NF-L in the serum samples collected along the duration of the study (Fig. 7) were not consistent with the histologic evidence of tissue destruction, with presence of myelin-laden CD68^+^ macrophages (Fig. 2a) and elevated pro-inflammatory cytokines and other factors (Fig. 6).

**Fig. 7.**
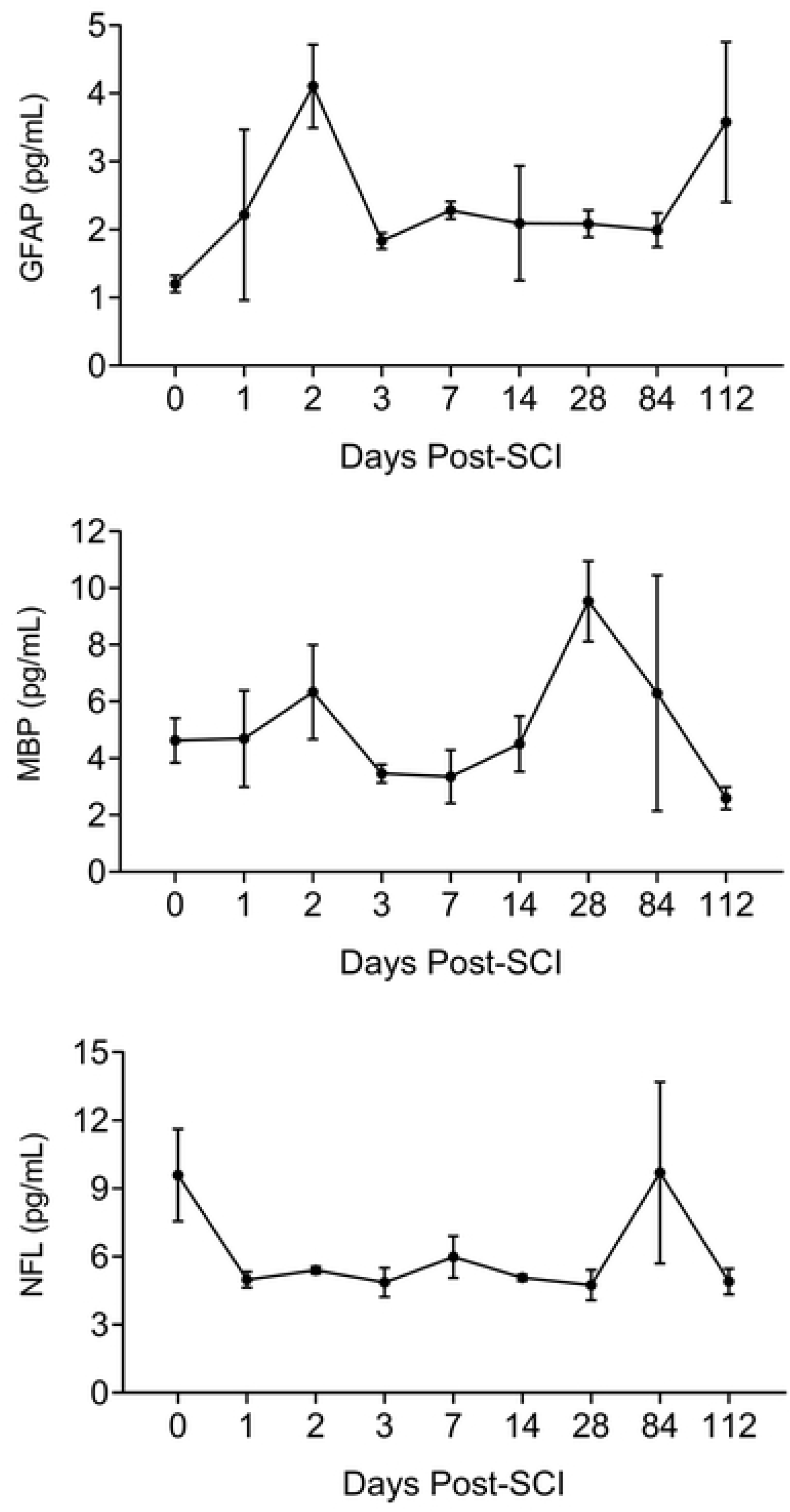
Biomarkers of spinal cord damage in the serum post-SCI.

## DISCUSSION

In this systematic study on the progression of SCI in the rat model we determined that a localized neurotrauma initiates two types of severe, destructive inflammatory response; the cavity of injury (COI) that progresses to a syrinx deep in the spinal cord and arachnoiditis at the surface that becomes a scar. The duration of both types of inflammatory response is extraordinarily long and is characterized by severe infiltration by phagocytizing CD68^+^/CD163^-^, M1 macrophages that continue to destroy myelin around the COI beyond week 16 post-SCI. Spinal cord reaction to severe inflammation is represented by progressively severe astrogliosis walling off the COI and arachnoiditis (Fig. 8). The anti-inflammatory activity deduced from the systematic histology of the sites of SCI involves; (1) the spinal cord-sparing enclosure of inflammation within the COI with concurrent sequestration of the fluid in the COI and astroglial reaction, (2) the gradual reduction of numbers of macrophages in the COI after week 4 post-SCI (Fig. 2a). Since arachnoiditis progresses from a granulomatous infiltration to a mature scar and does not contain astrocytes or other glial cells, we identify this as a distinct pathology different from the commonly described “glial scar” [36].

**Fig. 8.**
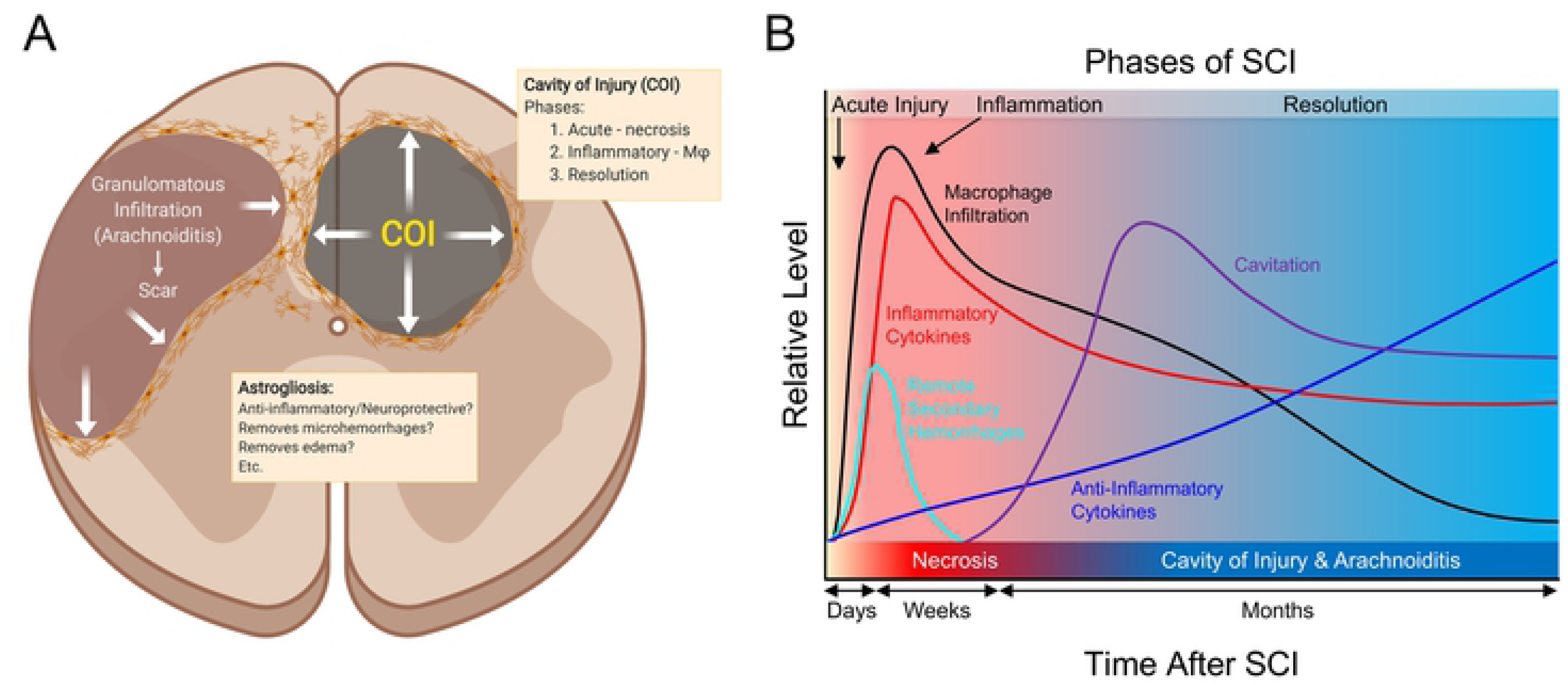
Mechanisms involved in spinal cord injury.

Spinal cord injury results in massive necrosis and hemorrhage where all cellular and vascular components of the tissue are destroyed. Areas of necrosis deep in the spinal cord and surrounded by the surviving spinal cord tissue become converted into the cavity of injury (COI) where inflammation is localized and fluid accumulates. The COI apparently expands during the long *Inflammatory Phase* and is surrounded and thus limited by gradually growing in the density wall of reactive astrogliosis while the severity of macrophage infiltration declines in the *Resolution Phase*. Areas of necrosis on the surface of the spinal cord are infiltrated by inflammatory granulomatous cells including macrophages, fibroblasts and capillary blood vessels from the subarachnoid space thus the name arachnoiditis. It appears to expand against the spinal cord tissue which recedes and forms a wall of gradually more intense astrogliosis. Over the time the area of arachnoiditis is fibrosed and becomes a typical scar devoid of glial cells therefore it is not a “glial scar” but rather a scar surrounded by astroglial reaction.

Analysis of T2 MR images indicates reduction of G/W border around the COI implying persistent vasogenic edema related to protracted inflammation, a pathologic mechanism with potential importance in the traumatic brain injury and stroke not previously considered. The severity of inflammation initiated after SCI is of a higher intensity and longer duration than is observed in trauma in an extra-neural soft tissue. This is related to the large quantity of damaged myelin, a potently immunogenic, pro-inflammatory material in the white matter injury in models of neurotrauma [38, 39] and of multiple sclerosis [40] and to a potential mechanism of a vicious cycle sustaining severe inflammation. A large amount of damaged myelin is associated with the initiation of chemotaxis of numerous macrophages into the necrotic site starting at day 3 post-SCI. In parallel with myelin-phagocytizing, activated macrophages high levels of pro-inflammatory factors were released in the spinal cord including pro-inflammatory cytokines; IFN-gamma, TNF-alpha, IL1-beta, IL-6 (Fig. 6), chemokines (CC and CXC), and reactive oxygen species [1]. The additional supply of damaged myelin likely sustained further macrophage chemotaxis [38, 41]. Given the mechanism proposed here for the vicious cycle of damage and inflammation, the extraordinary duration of the inflammation, beyond 16 weeks post-SCI can be explained. The prevalence of CD68^+^/CD163^-^, RBC- and myelin-phagocytizing macrophages in the COI and surrounding the COI is indicative of the M1 phenotype, a presumed pro-inflammatory macrophage [40] and supports the notion that severe inflammation contributes to progressive spinal cord destruction.

A dual anti-CD68/anti-CD163 antibody labeling of paraffin-embedded spinal cord tissues was performed since these epitopes are considered reliable indicators of M1 and M2 macrophage phenotypes, respectively, *in vivo* [42]. The polarization of macrophages participating in inflammation [26] - from an inflammatory M1 phenotype, induced by TNF-alpha, IFN-gamma and IL-6 [26] into an anti-inflammatory M2 phenotype, induced by IL-10 [43] is not evident in the COI along the course of the *Inflammatory Phase* of the SCI. Progressive inhibition of inflammation in the COI is not likely due to an immune process alone. It appears more likely that a spinal cord reaction markedly contributed to this inhibition. Such anti-inflammatory activity was found to be potent in the SCI of the Long Evans Shaker (LES) rat [15] where severe astrogliosis forms in response to lack of myelin [20]. The dynamic changes in the population of CD68/CD163-labeled monocyte progenitors of the bone marrow (Fig. 3b) are also suggestive that activity inhibiting the inflammation in the COI, does not involve increased levels of M2 type macrophages.

Although the process of expansion of the COI and of arachnoiditis with the concurrent destruction of the adjacent spinal cord is not analyzed directly in this study it is implied (Fig. 1A; 2a) from the continuous presence of CD68^+^/CD163^-^ pro-inflammatory macrophages containing myelin granules beyond 16 weeks post-SCI. Progressive expansion of the volume of the COI following SCI has been previously indicated [34, 44]. It appears that an anti-inflammatory/neuroprotective therapy for SCI should involve reducing the numbers of macrophages during the inflammatory phase to inhibit myelin damage (Fig. 2b) [24].

Astrogliosis is a hallmark of an integrated tissue reaction that results in the containment of a severe inflammation initiated by the SCI, its inhibition and ultimately elimination (Fig. 4). Although a variety of neuroprotective functions attributed to astrogliosis have been proposed [3] the indication of its negative effect, inhibition of the axonal regeneration by virtue of a “glial scar” barrier persists [1]. Herein we propose that in the two types of inflammatory reaction to neurotrauma; the COI and arachnoiditis, axons do not enter the liquid environment of COI since they don’t swim and require a bridge to cross COI [15]. In arachnoiditis the granulomatous tissue that progressively matures into a scar, does not contain glial cells (Fig. 3a) therefore it ceases to be a part of the CNS and is an unsuitable milieu for axonal regeneration. Both COI and arachnoiditis are walled off by increasingly severe astrogliosis from the reminder of the spinal cord. Reactive astrocytes however, apparently do not inhibit axonal regeneration [15,45,46].

Removal of RBCs from the tissue surrounding the necrosis was associated with GFAP^+^ stellate cells interpreted as reactive astrocytes and not by CD68^+^ macrophages. Although reactive astrocytes have been shown to be phagocytic in penumbra of ischemic brain [47], the *astrocytic erythrophagocytosis* has not been previously reported and is considered a protective mechanism where scattered RBCs are rapidly removed from the spinal cord to avoid macrophage chemotaxis leading to their activation and tissue damage. The astrocytic erythrophagocytosis is a histologically obvious process that may constitute a pathologic basis of limited neurotrauma that is not detected on computed tomography and MRI such as in concussion. Astrogliosis around the acute injury and the forming COI may be related to restoration of homeostasis [2,28,29] via a number of distinguishable mechanisms including inhibition of inflammation, erythrophagocytosis and removal of tissue edema [33, 48].

The ependymal reaction to injury and inflammation post-SCI involves proliferation, migration and transient differentiation into astrocytic lineage (Fig. 5b). The mechanisms behind the plasticity of ependymal cells and their migration [49] appear to be a response to the mechanical and inflammatory damage and can result in trans-differentiation of ependymal cells into astrocytes [49], oligodendrocytes 50 and neural progenitors [51]. The importance of this plasticity includes an active support and guidance provided to regenerating axons [15,36,52].

Although the proteomic data support the notion of severe inflammation, consistent with persistence of CD68^+^/CD163^-^ M1 macrophages, the decline of their numbers combined with the reduction in the levels of pro-inflammatory cytokines indicates a progressive increase in the anti-inflammatory activity. In prior studies the levels of inflammatory cytokines and other factors have been measured at acute stages in the spinal cord, CSF or serum of SCI animal models and SCI patients [8, 37] with studies of long progression of inflammation lacking. Remarkable elevation of TNF-alpha and IL-1beta was associated with increased activity of astrocytes and microglia and infiltrating leukocytes [8]. Pro-inflammatory cytokines secreted by reactive astrocytes, microglia and infiltrating macrophages have been shown to injure myelin sheaths and axons in the spinal cord surrounding the trauma [31, 53]. In notable exceptions; CX3CL1 (fractalkine), CD86, and IL-13 show consistent gradual increase in levels of the course of the SCI in the present study, and may have been related to tissue anti-inflammatory/neuroprotective activity. Increase in fractalkine activity has been proposed to be neuroprotective [46] but not considered beyond the acute phase in neurotrauma. Elevation of fractalkine in astrocytes resulted in increased expression of CX3CR1 in co-cultured microglia resulted in an anti-inflammatory effect [54] suggesting that elevated fractalkine may correlate with astrogliosis. Elevated IL-13, IL-4 and IL-10 have been shown to induce polarization of macrophage infiltration in the SCI to the M2 type considered anti-inflammatory and neuroprotective [55]. Relatively low levels of these 3 cytokines with elevation of pro-inflammatory cytokines including IL-1beta, IFN-gamma and IL-6 with consistent predominance of CD68^+^/CD163^-^ macrophages that actively phagocytized myelin in the COI support the notion of the lack of polarization of M1 to M2 macrophages during the *Inflammatory Phase* and beyond 4 months post-SCI. Therefore, the inhibition and elimination of inflammation during the *Resolution Phase* may be related to a novel mechanism involving a spinal cord anti-inflammatory activity, specifically to astrogliosis.

## ACKNOWLEDGEMENTS

Funding support provided in part by the Medical University of Lublin. The authors wish to acknowledge excellent technical support of Mary Jo Smith and Mary Bruni for histologic and immunohistochemical preparations.

